# Global gene expression patterns in response to white patch syndrome: Disentangling symbiont loss from the thermal stress response in reef-building coral

**DOI:** 10.1101/2019.12.13.875989

**Authors:** Carly D. Kenkel, Veronique J.L. Mocellin, Line K. Bay

**Affiliations:** Department of Biological Sciences, University of Southern California, 3616 Trousdale Parkway, Los Angeles, CA 90089, USA; Australian Institute of Marine Science, PMB No. 3, Townsville MC, QLD 4810, Australia

**Keywords:** *Porites lobata*, bleaching, symbiosis, meta-analysis, Tag-Seq

## Abstract

The mechanisms resulting in the breakdown of the coral symbiosis once the process of bleaching has been initiated remain unclear. Distinguishing symbiont loss from the abiotic stress response may shed light on the cellular and molecular pathways involved in each process. This study examined physiological changes and global gene expression patterns associated with white patch syndrome (WPS) in *P. lobata*, which manifests in localized bleaching independent of thermal stress. In addition, a meta-analysis of global gene expression studies in other corals and anemones was used to contrast differential regulation as a result of abiotic stress from expression patterns correlated with symbiotic state. Symbiont density, chlorophyll *a* content, holobiont productivity, instant calcification rate, and total host protein content were uniformly reduced in WPS relative to healthy tissue. While expression patterns associated with WPS were secondary to fixed effects of source colony, specific functional enrichments suggest that the viral infection putatively giving rise to this condition affects symbiont rather than host cells. The meta-analysis revealed that expression patterns in WPS-affected tissues were significantly correlated with prior studies examining short-term thermal stress responses. This correlation was independent of symbiotic state, as the strongest correlations were found between WPS adults and both symbiotic adult and aposymbiotic coral larvae experiencing thermal stress, suggesting that the majority of expression changes reflect a non-specific stress response. Across studies, the magnitude and direction of expression change among particular functional enrichments suggests unique responses to stressor duration, and highlights unique responses to bleaching in an anemone model which engages in a non-obligate symbiosis.

## Introduction

Dinoflagellates in the family Symbiodiniaceae are obligate endosymbionts of reef-building corals. Photosynthetically-derived nutrition provided by symbionts largely fuels the process of coral host calcification which builds the three dimensional reef structure that supports ecosystem function (C. A. Oakley & Davy, 2018; Rogers, Blanchard, & Mumby, 2014). Host energetic status, calcification, and consequently, ecosystem function are impaired in the process known as ‘coral bleaching’ in which abiotic stress leads to the breakdown of the symbiosis. Thermal stress events are the most common cause of mass bleaching worldwide and are increasing in frequency and severity as a result of global climate change (T. P. Hughes et al., 2018). Resolving the mechanisms that facilitate initiation and maintenance of a healthy symbiosis, as well as those underpinning its breakdown will be essential for developing solutions to the ongoing coral reef crisis (Weis, 2019).

A number of biological processes are thought to be involved in mediating the establishment and long-term maintenance of the coral symbiosis including host-microbe signaling, regulation of the host innate immune response and cell cycle, phagocytosis, and cytoskeletal rearrangement (Davy, Allemand, & Weis, 2012). Five different bleaching mechanisms have been observed and/or proposed, but it is unknown which mechanism(s) predominate in natural bleaching events, whether mechanisms differ among species, and to what extent they may interact (reviewed in (C. A. Oakley & Davy, 2018; Weis, 2008)). A comparative study which exposed clonal replicates of *Aiptasia pallida* to a combination of controlled abiotic stressors found that expulsion of intact symbionts from host cells was the predominant response to heat and/or light stress, while *in situ* degradation of symbiont cells and host-cell detachment also played a role in the bleaching response to acute cold shock (Bieri, Onishi, Xiang, Grossman, & Pringle, 2016). An earlier study, however, reported that host cell detachment was the predominant mechanism of bleaching in response to elevated temperature stress (Gates, Baghdasarian, & Muscatine, 1992), whereas *in situ* degradation of symbiont cells was the primary response of a temperate coral to annual thermal bleaching (Ainsworth & Hoegh-Guldberg, 2008). It is possible that these mechanisms may occur in combination depending on the magnitude and duration of the stress (Bieri et al., 2016; Gates et al., 1992), but additional work is needed to quantify their relative importance (C. A. Oakley & Davy, 2018).

High-throughput transcriptomic, proteomic, and metabolomic approaches have provided additional mechanistic insight into the cellular functions which are disrupted as a result of thermal stress and bleaching. Unfolded protein and oxidative stress response pathways have been universally highlighted (Barshis et al., 2013; Cziesielski Maha J. et al., 2018; Clinton A. Oakley et al., 2016). Transcriptomic and proteomic studies have also repeatedly identified changes to cytoskeleton and extracellular matrix proteins (DeSalvo, Sunagawa, & Voolstra, 2010; DeSalvo et al., 2008; Clinton A. Oakley et al., 2016), which has been interpreted as evidence supporting the role of autophagy and apoptosis in the mechanism of bleaching (C. A. Oakley & Davy, 2018). However, other investigations into thermal stress responses of aposymbiotic juvenile coral life history stages have also identified differential regulation of cytoskeletal components and heat shock proteins (Meyer, Aglyamova, & Matz, 2011; Polato et al., 2010; Rodriguez-Lanetty, Harii, & Hoegh-Guldberg, 2009; Voolstra, Schnetzer, et al., 2009), suggesting that these responses are not necessarily unique to the process of bleaching, but rather reflect a more general cellular stress response common to all organisms (Kültz, 2005). Consequently, disentangling the thermal stress response from the process of symbiont loss may help distinguish among the prospective cellular and molecular mechanisms responsible for bleaching.

The coral symbiosis can break down in response to a variety of stressors, both biotic and abiotic (Brown & Howard, 1985); however, few studies have examined molecular or cellular responses to bleaching in non-heat related contexts. Pharmacological agents can be used to induce bleaching independent of temperature, and indeed, host cell detachment has been observed in response to chemical stress (Sawyer & Muscatine, 2001). It is also possible to bleach facultatively symbiotic *Aiptasia pallida* anemones using menthol without impairing their ability to later re-establish symbiosis (Matthews et al., 2016). A proteomic comparison of symbiotic and aposymbiotic anemones generated via menthol bleaching identified an increase in proteins involved in mediating reactive oxygen stress (Clinton A. Oakley et al., 2016). Oxidative stress response pathways were also differentially regulated in menthol-bleached *Aiptasia* re-infected with a heterologous symbiont type (Matthews et al., 2017), suggesting that this pathway responds, at least in part, to symbiotic state independent of ambient temperature. While some progress has been made in anemone models, to our knowledge, similar attempts to disentangle bleaching from thermal stress have yet to be undertaken in reef-building corals.

This study aimed to fill this gap by taking advantage of a coral disease which does not harm host tissue integrity, but results in localized bleaching independent of thermal stress. White patch syndrome (WPS) affects massive poritids in the Indo-Pacific (Lawrence, Davy, Wilson, Hoegh-Guldberg, & Davy, 2015; Raymundo, Rosell, Reboton, & Kaczmarsky, 2005; Roff, Ulstrup, Fine, Ralph, & Hoegh-Guldberg, 2008). In corals affected by WPS, small patches of bleached tissue are present without any necrosis or apparent impact to host tissues (Lawrence et al., 2015; Roff et al., 2008). A comparative analysis of symbiont photosynthetic function in response to different coral diseases found that WPS was the only syndrome to impact symbiont photosynthetic function in a pattern similar to thermal stress induced bleaching (Roff et al., 2008). Additional physiological data revealed that patches contained significantly reduced chlorophyll *a* content, but that soluble host protein content was similar to healthy tissue, suggesting that WPS primarily impacts photosymbionts (Lawrence et al., 2015). Transmission electron microscopy images generated in the same study confirmed that while host tissue remained intact within patches, there was a significant increase in virus-like particles in both coral and Symbiodiniaceae cells, suggesting that the causative agent of WPS may be a virus which infects either the host or symbiont (Lawrence et al., 2015).

We used global gene expression analyses in combination with a suite of ecophysiological assays to compare the molecular and physiological responses of healthy and WPS tissue patches within and among eight independent coral colonies. In addition, we conducted a meta-analysis to compare and contrast WPS expression responses with prior studies examining expression responses to symbiotic state (Matthews et al., 2016), short-term thermal stress in adult and aposymbiotic larval corals (Dixon et al., 2015; Meyer et al., 2011), and long-term thermal stress in adult corals (Davies, Marchetti, Ries, & Castillo, 2016). We find that all aspects of coral holobiont physiology are negatively impacted by WPS, but that global gene expression patterns associated with WPS are secondary to fixed effects of source colony. Overall, WPS expression patterns are most similar to short-term thermal stress responses regardless of symbiotic status, and specific enrichments suggest that if WPS is caused by a viral infection, it does not significantly impact *Porites* host tissue.

## Methods

### Sample Collection

Eight colonies of *Porites lobata* were sampled between 19 - 21st April 2015 from the northern side of Pandora reef (18.813878° S, 146.432784° E) under Great Barrier Reef Marine Park Authority permit G12/35236.1 and G14/37318.1. Colonies were haphazardly selected based on exhibiting the WPS phenotype. Four cores were taken from each of 8 corals that exhibited WPS symptoms: (1) two from the center of an affected patch and (2) two from an adjacent area of normal tissue using an underwater drill (18V Nemo Power Tools, USA) fitted with a custom-built 2-cm diameter coring attachment with diamond tip teeth. One core of each origin was snap frozen in liquid nitrogen after collection for RNA analysis. The second core was first photographed and assessed for photosynthesis and respiration capability, before being snap frozen for further physiological analyses in equal conditions, approximately 6 hours after collection.

### Physiological Assays

Net photosynthesis and respiration were assayed on the day of collection following the two-point method originally described and validated for *A. millepora* (Strahl et al., 2015). Cores were incubated for 100 min at either 250 μmol/photon/m^−2^/s^−1^ (net photosynthesis) or full darkness (respiration). During incubations each core was enclosed in a 600-ml acrylic chamber placed onto custom built tables with rotating magnets, which served to power stir bars within each chamber to facilitate water mixing. To account for potential changes in oxygen content due to the metabolic activity of other microorganisms in the seawater, two chambers without corals were used as blanks in each run. At the end of the incubation period the O_2_ concentration of the seawater in each chamber was measured using a hand-held dissolved oxygen meter (HQ30d, equipped with LDO101 IntelliCAL oxygen probe, Hach, USA). Values from blank chambers were subtracted from measures made in coral chambers and the subsequent rate of net photosynthesis was related to coral surface area, calculated in μg O_2_/cm^2^/min. Gross photosynthesis was calculated as the sum of net photosynthesis and respiration. Holobiont productivity (P:R) was quantified as the ratio of oxygen production (gross photosynthesis) to oxygen consumption through respiration.

To determine total alkalinity, a 120 ml subsample of seawater from each incubation chamber was fixed immediately following the light incubation with 0.5 mg mercuric chloride (final concentration 4 mg/L). Light calcification rates were determined from changes in alkalinity quantified with the alkalinity anomaly technique (Chisholm & Gattuso, 1991) using a Titrando 855 Robotic Titrosampler (Metrohm AG, Switzerland). Values from blank chambers were subtracted from measures made in coral chambers and calcification rate was related to coral surface area and calculated in μM CaCO_3_/cm^2^/min.

To quantify symbiont density, photosynthetic pigments and protein content, tissues were removed from host skeletons using an air gun and 0.2 μM filtered seawater (FSW) and homogenized for 60 s using a Pro250 homogenizer (Perth Scientific Equipment, AUS). Coral skeletons were rinsed with 5% bleach then dried at room temperature. Skeletal surface area was quantified using the single wax dipping method (Stimson & Kinzie, 1991) and skeletal volume was determined by calculating water displacement in a graduated cylinder. A 200-μl aliquot of tissue homogenate was fixed with equal volume of 10% formalin in FSW and used to quantify *Symbiodiniaceae* cell density. The average cell number was obtained from four replicate haemocytometer counts of a 1-mm^3^ area and cell density was related to coral surface area and expressed as cells/cm^2^. A 1 mL aliquot of the tissue homogenate was centrifuged for 3 minutes at 1500 × *g* at 4°C and the pellet was stored at −80°C for chlorophyll analysis. The remaining homogenate was separated into host and symbiont fractions by cold centrifugation (4°C) for 10 min at 3200 × g. To quantify chlorophyll concentrations, algal pellets were resuspended in 1 mL chilled 95% Ethanol. The homogenate was sonicated on ice for 30 seconds at 40% amplitude and centrifuged for 5 minutes at 10,000 ×*g* at 4°C. A 200 μL aliquot of sample extract was loaded to a 96-well plate, and absorbance was recorded at 632, 649, and 665 nm. Chlorophyll *a* concentration was calculated with the equation in (Ritchie, 2008), normalized to surface area, and expressed as μg/cm^2^. A commercial colorimetric protein assay kit (DCTM Protein Assay Kit, Bio-Rad, Hercules, USA) was used to quantify total protein content of the coral host tissue. A 50 μL aliquot of *Symbiodiniaceae*-free coral tissue sample was digested using 50 μL 1M Sodium hydroxide for 1 hour at 90°C. The plate was centrifuged for 3 minutes at 1500 × *g*. Following manufacturer’s instructions, 5 μL of digested tissue was mixed with 25 μL alkaline copper tartrate solution and 200 μL dilute Folin reagent in a 96-well plate. Absorbance at 750 nm was recorded after a 15-minute incubation. Sample protein concentrations were calculated using a standard curve of bovine serum albumin ranging from 0 and 2000 μg mL^−1^. Protein concentrations were normalized to total homogenate volume and coral surface area, and expressed as mg/cm^2^.

### Tag-Seq Library Preparation

Total RNA was extracted using an Aurum Total RNA mini kit (Bio-Rad, CA), with minor modifications. Briefly, samples homogenized in lysis buffer were kept on ice for one hour with occasional vortexing to increase RNA yields, which was followed by centrifugation for 2 minutes at 14000 × g to precipitate skeleton fragments and other insoluble debris; 700 μl of the supernatant was used for RNA purification. At the final elution step, the same 25 μl of elution buffer was passed twice through the spin column to maximize both yield and concentration of eluted RNA. Samples were DNAse treated as in (Kenkel et al., 2011). One μg of total RNA per sample was used for tag-based RNA-seq, or Tag-Seq (Meyer et al., 2011), with modifications for sequencing on the Illumina platform (the current versions of the laboratory and bioinformatic protocols for Tag-Seq are maintained at https://github.com/ckenkel/tag-based_RNAseq). Tag-Seq has been shown to generate more accurate estimates of protein-coding transcript abundances than standard RNA-seq, at a fraction of the cost (Lohman, Weber, & Bolnick, 2016).

### Reference Transcriptome sequencing, assembly and annotation

To generate a *P. lobata* reference transcriptome, 5 replicate fragments of a single coral colony were subject to a two-week temperature stress experiment as described in (Kenkel & Bay, 2018). Snap frozen samples from control (27°C, days 4 and 17) and heat (31°C, days 2, 4 and 17) treatments were crushed in liquid nitrogen and total RNA was extracted using an Aurum Total RNA mini kit (Bio-Rad, CA). RNA quality and quantity were assessed using the NanoDrop ND-200 UV-Vis Spectrophotometer (Thermo Scientific, MA) and gel electrophoresis. RNA samples from replicate fragments were pooled in equal proportions and 1.1 ug was shipped on dry ice to the Oklahoma Medical Research Foundation NGS Core where Illumina TruSeq Stranded libraries were prepared and sequenced on one lane of the Illumina Hiseq 3000/4000 to generate 2 × 150 PE reads.

Sequencing yielded 76 million raw PE reads. The *fastx_toolkit* (http://hannonlab.cshl.edu/fastx_toolkit) was used to discard reads < 50 bp or having a homopolymer run of ‘A’ ≥ 9 bases, retain reads with a PHRED quality of at least 20 over 80% of the read and to trim TruSeq sequencing adaptors. PCR duplicates were then removed using a custom perl script (https://github.com/ckenkel/annotatingTranscriptomes). Remaining high quality filtered reads (19 million paired reads; 3 million unpaired reads) were assembled using Trinity v 2.0.6 (Grabherr et al., 2011) using the default parameters and an *in silico* read normalization step at the Texas Advanced Computing Center (TACC) at the University of Texas at Austin.

The holobiont assembly was filtered to produce a host-specific assembly using a series of hierarchical BLAST searches against the *Acropora digitifera* (Shinzato et al., 2011) and *Symbiodinium kawagutii* (S. Lin et al., 2015) proteomes, and NCBI’s nr database (Pruitt, Tatusova, & Maglott, 2005) following (Kenkel & Bay, 2017). Annotation was performed following the protocols and scripts described at https://github.com/ckenkel/annotatingTranscriptomes. Contigs were assigned putative gene names and gene ontologies using a BLASTx search (E value <= 10^−4^) against the UniProt Knowledgebase Swiss-Prot database (UniProt Consortium, 2015). KOG (EuKaryotic Orthologous Groups) annotations were computed using eggnog-mapper (Huerta-Cepas et al., 2017) based on eggNOG 4.5 orthology data (Huerta-Cepas et al., 2016) and KEGG (Kyoto Encyclopedia of Genes and Genomes) id’s using the KAAS server (http://www.genome.jp/kegg/kaas/ (Moriya, Itoh, Okuda, Yoshizawa, & Kanehisa, 2007). The stats.sh command of the BBMap package (Bushnell, 2014) was used to calculate GC content of host transcriptomes. Transcriptome completeness was evaluated through comparison to the Benchmarking Universal Single-Copy Ortholog (BUSCO v2) (Simão, Waterhouse, Ioannidis, Kriventseva, & Zdobnov, 2015) set using the gVolante server (https://gvolante.riken.jp/analysis.html).

### Tag-Seq Sequencing

Sixteen Tag-Seq libraries were sequenced across four lanes of the HiSeq 2500 at the University of Texas at Austin Genome Sequencing and Analysis Facility. Resulting reads were concatenated by sample before processing. On average 7.3 million sequences were generated per library (median: 5.3 million, range: 0.2 - 43.8 million), for a total of 117.7 million raw reads. Custom perl scripts were used to discard non-template sequence reads (defined as reads lacking the appropriate sequencing primer at the 5’ end) while also trimming sequencing primers. The *fastx_toolkit* (http://hannonlab.cshl.edu/fastx_toolkit/index.html) and BBTools (https://sourceforge.net/projects/bbmap/) were used to further remove reads shorter than 20 bases or exhibiting homopolymer runs of ‘A’ in excess of 8 bases or those exhibiting significant matches to illumina indexes. From these, only reads with PHRED quality of at least 20 over 70% of the sequence were retained. See Table S1 for a summary of reads retained in each filtering step.

SHRiMP2 (David, Dzamba, Lister, Ilie, & Brudno, 2011) was used to map filtered reads against the *P. lobata* host reference concatenated to the *Cladocopium* strain C15 reference transcriptome (Shinzato, Inoue, & Kusakabe, 2014) as in (Kenkel, Moya, Strahl, Humphrey, & Bay, 2018). Resulting read counts were summed by isogroup (groups of sequences putatively originating from the same gene, hereafter referred to as genes), sample, and focal organism (host, symbiont). On average, 2.1 million reads per sample were mapped to 25,222 genes in the host transcriptome, while 0.7 million reads per sample were mapped to 10,629 genes in the symbiont transcriptome. Due to the low number of reads mapping to the symbiont reference, only host expression data were statistically analyzed.

### Statistical analyses

All statistical analyses were completed in R 3.6.0 (R Core Team & Others, 2017) and scripts and input files can be found at https://github.com/ckenkel/PoritesWPS. For the physiological data, we modelled the fixed effect of condition (levels: healthy, WPS) on *Symbiodiniaceae* cell density, chlorophyll a content, gross photosynthesis, total host protein and holobiont instant calcification rate using the *lme* command of the *nlme* package (Pinheiro, Bates, DebRoy, Sarkar, & Others, 2014). Source coral colony identity was included in all models as a scalar random factor. Models were assessed for normality of residuals and homoscedasticity. The *prcomp* command of the *stats* package was used to conduct a principal components analysis of the host physiology data and the first principal component was used as an additional trait for correlation with gene expression.

For gene expression counts, the package *arrayQualityMetrics* (Kauffmann, Gentleman, & Huber, 2009) was used to identify outliers. Both the ‘normal’ and ‘WPS’ samples from coral genet 4 were identified as outliers and excluded from further analysis in the host dataset, likely due to the poor quality of sample 4N (the healthy sample, Table S1). Statistical analyses were conducted using the package *DESeq* (Anders & Huber, 2010). A Cox-Reid adjusted profile likelihood model specifying coral colony of origin and phenotype (normal or WPS) was used to obtain the maximum dispersion estimate of the raw counts data. The bottom 10% quantile of genes was identified as the statistic which best satisfied the assumptions of independent filtering as implemented in *genefilter* (Gentleman, Carey, Huber, & Hahne, 2019) and these low-expression genes were removed from each dataset prior to formal hypothesis testing. This left 22,630 highly expressed genes. A series of generalized linear models implemented using the function *fitnbinomGLMs* were used to test the effects of WPS phenotype and PC1 of the physiology data on the expression of individual genes while controlling for fixed effects of colony of origin. The *KOGMWU* package (Dixon et al., 2015) was used to test for rank-based enrichment of euKaryotic Orthologous Groups (KOG) terms among differentially expressed genes by WPS phenotype. The GO_MWU package (Wright, Aglyamova, Meyer, & Matz, 2015) was used to test for rank-based enrichment of gene ontology terms. Both enrichment tests were conducted using signed log P-values as the measure of interest as in (Strader, Aglyamova, & Matz, 2016).

### Meta-analysis of gene expression response to bleaching and thermal stress

The *KOGMWU* package was used to compare KOG term enrichments identified for the WPS phenotype with existing cnidarian datasets which quantified global gene expression responses to thermal stress and symbiotic state to distinguish molecular responses specifically determined by symbiotic state. Comparative studies included short-term thermal stress responses in aposymbiotic *Acropora millepora* larvae (5 days (Meyer et al., 2011)), and symbiotic adult *A. millepora* (3 days, (Dixon et al., 2015)), long-term thermal stress response in symbiotic adult *Siderastrea siderea* (95 days, (Davies et al., 2016)), and symbiotic state independent of thermal stress in the model anemone *Aiptasia pallida* (Matthews et al., 2017). For the *S. siderea* and *A. pallida* datasets, tables of mapped read counts and reference transcriptomes and in the case of *A. pallida*, statistical R scripts, were obtained from the respective corresponding authors. *DESeq* (Anders & Huber, 2010) was used to re-analyze both datasets. The *A. pallida* re-analysis largely followed statistical methods originally described in Matthews et al. (2017) save that the categorical comparison of interest was aposymbiotic vs symbiotic anemones (as opposed to the type of symbiont hosted). The *S. siderea* re-analysis reduced the dataset to the ambient vs. elevated temperature comparison only. The *A. millepora* datasets are included as example data with the *KOGMWU* package. For all supplementary datasets, KOG annotations were updated using eggnog-mapper (Huerta-Cepas et al., 2017) based on eggNOG 4.5 orthology data (Huerta-Cepas et al., 2016). Pearson correlations were calculated to evaluate similarity among up- and down-regulated KOG classes across datasets.

## Results

### White patch syndrome physiology

All physiological parameters were negatively impacted within WPS affected tissues. Symbiont density decreased by 1.5 × 10^6^ cells/cm^2^ on average in WPS infected tissue relative to paired healthy controls (P = 0.001, Fig. 1a). Similarly, chlorophyll *a* content declined by 18.4 μg/cm^2^ (P < 0.001, Fig. 1b) and P:R ratios were reduced by 0.91 on average (P < 0.001, Fig. 1c). Soluble host protein content was 256.7 mg/cm^2^ lower (P = 0.046, Fig. 1d) and calcification rate decreased by 11.7 μmol CaCO_3_/cm^2^/min in WPS affected tissue compared to healthy tissue on average. A principal components analysis revealed that overall, nearly 60% of the variation in the physiological data was driven by WPS phenotype (Fig. S1a) whereas source colony effects were secondary (Fig. S1b).

**Figure 1.**
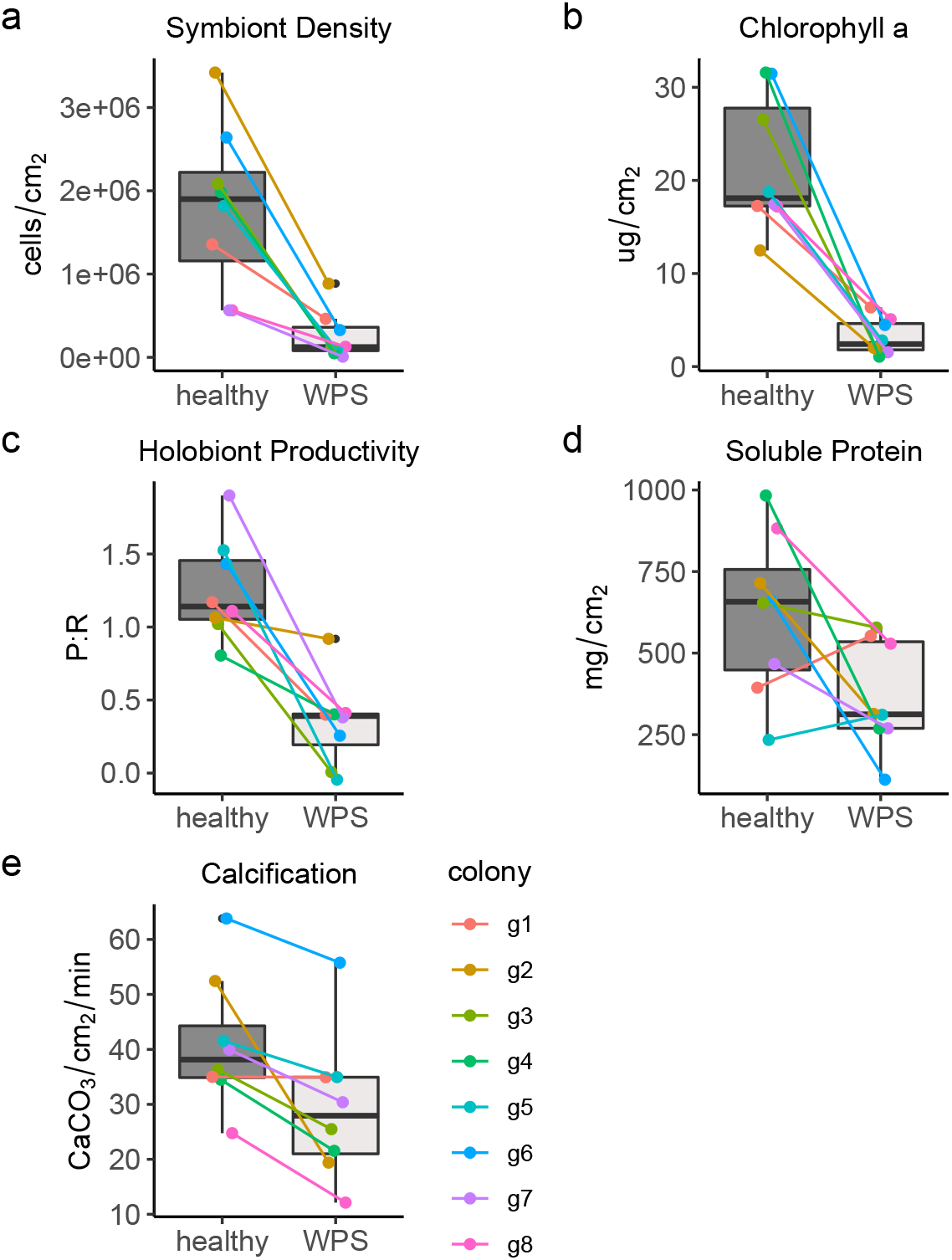
Physiological responses to white patch syndrome (WPS). Boxplots show the distribution of traits by phenotype. The median is indicated by the thick black line and the box represents the range between the upper and lower quartile. Whiskers extend maximally 1.5 times beyond the inter-quartile range and outliers are indicated by black circles. Colored lines indicate paired samples originating from each unique source colony.

### Porites lobata *reference transcriptome*

The initial holobiont assembly contained 125,405 contigs over 400 bp in length (N_50_ = 1537). Of these, 52 were discarded as matching non-mRNAs (12 rRNA, 40 mitochondrial). Following screening for biological contamination, 42,793 contigs had a best match to the *Acropora digitifera* proteome, and of these, 39,472 matched either a metazoan or had no match in NCBI’s nr database. An additional 3,243 contigs matched neither proteome, but exhibited a best hit to a Cnidarian in the nr database and were also retained. This *P. lobata* specific 42,715 contig assembly represented 23,349 isogroups (~genes) with an average length of 1,598 bp and an N50 of 2,147). GC content of the host-specific assembly was 42%, consistent with other anthozoan transcriptomes where *Symbiodiniaceae* reads have been effectively filtered (Kenkel & Bay, 2017; Z. Lin et al., 2017). Protein coverage exceeded 0.75 for 43% of contigs. Comparison of this assembly to the core eukaryotic 248-gene set (Parra, Bradnam, & Korf, 2007) revealed 90.7% of KOGs were represented. Of the 978 core BUSCO gene set for metazoans (Simão et al., 2015), 89.57% were found to be complete, while an additional 2.86% were partially assembled indicating that the assembly is comprehensive, especially in comparison to other currently available resources for this species (Quek & Huang, 2019).

### Host expression associated with WPS

Unlike the physiological data, the largest effect detected in the expression dataset was source colony identity. Nearly 43% of genes in the host transcriptome (9,705) were significantly differentially expressed by source colony identity, whereas only 0.7% (148 genes) were differentially expressed by WPS phenotype alone (P_adj_<0.1, Fig. S2). An additional 396 genes (1.7%) showed an effect of both source colony and WPS. No genes exhibited significant correlations with the first principal component of the physiological data alone (‘PC1’, Fig. S2), but 10 genes showed a significant effect of WPS and PC1, while 31 genes were differentially expressed by source colony, WPS and PC1. A principal components analysis of the top most significantly differentially expressed genes confirmed that source colony identity was the major driver of variation in expression (PC1), but PC2 largely differentiated samples based on WPS phenotype, indicating that this condition is still a reasonably strong driver of host expression (Fig. S3). Of all the 585 genes exhibiting differential expression by WPS phenotype the direction of regulation was fairly evenly split, with 323 genes downregulated and 255 upregulated.

Rank-based functional enrichment analyses of expression changes associated with WPS highlight a general downregulation of mitochondrial metabolism, and upregulation of both cytoskeletal components and genes involved in RNA processing and chromatin organization. Significant gene ontology (GO) enrichments were detected for ‘biological process’ and ‘cellular component’ terms (Fig 2). The strongest enrichments were among cellular component terms, with ‘cytoskeletal part’ showing the most significant upregulation (GO:0044430, P_adj_< 0.001) and ‘mitochondrial part’ exhibiting the strongest downregulation (GO:0044429, P_adj_< 0.001, Fig. 2b). Biological process terms were less strongly regulated, but ‘chromatin organization’ topped upregulated terms (GO:0006325, P_adj_< 0.001) whereas ‘regulation of ossification’ was the most significant enrichment among downregulated genes (GO:0030278, P_adj_< 0.01). Similarly, the top three eukaryotic orthologous group (KOG) enrichments included ‘energy production and conversion’ (P_adj_< 0.001), ‘cytoskeleton’ (P_adj_< 0.001) and ‘RNA processing and modification’ (P_adj_< 0.005, Fig. 3). Differentially expressed genes annotated with the energy production and conversion term largely consisted of cytochrome oxidase subunits and other enzymes in the citric acid cycle and exhibited patterns of downregulation in WPS-affected tissues (Fig. S4) consistent with GO term enrichments.

**Figure 2.**
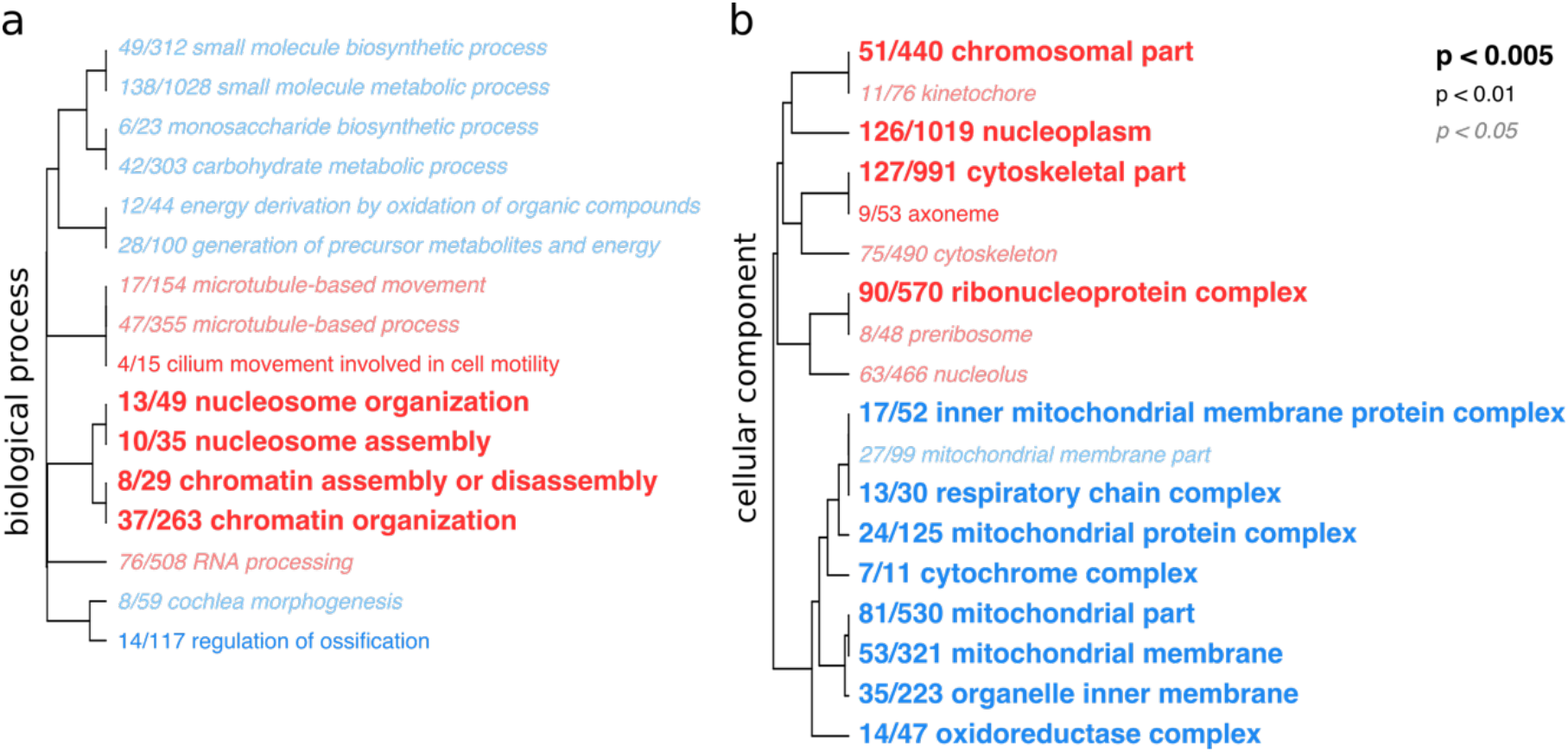
Hierarchical clustering of enriched (a) “biological process” and (b) “cellular component” gene ontology terms among generally upregulated (red) and downregulated (blue) genes in the coral host with respect to WPS phenotype. Font indicates the level of FDR-adjusted statistical significance. Term names are preceded by a fraction indicating the number of individual genes within each term that are differentially regulated with respect to WPS phenotype (unadjusted P < 0.05)

**Figure 3.**
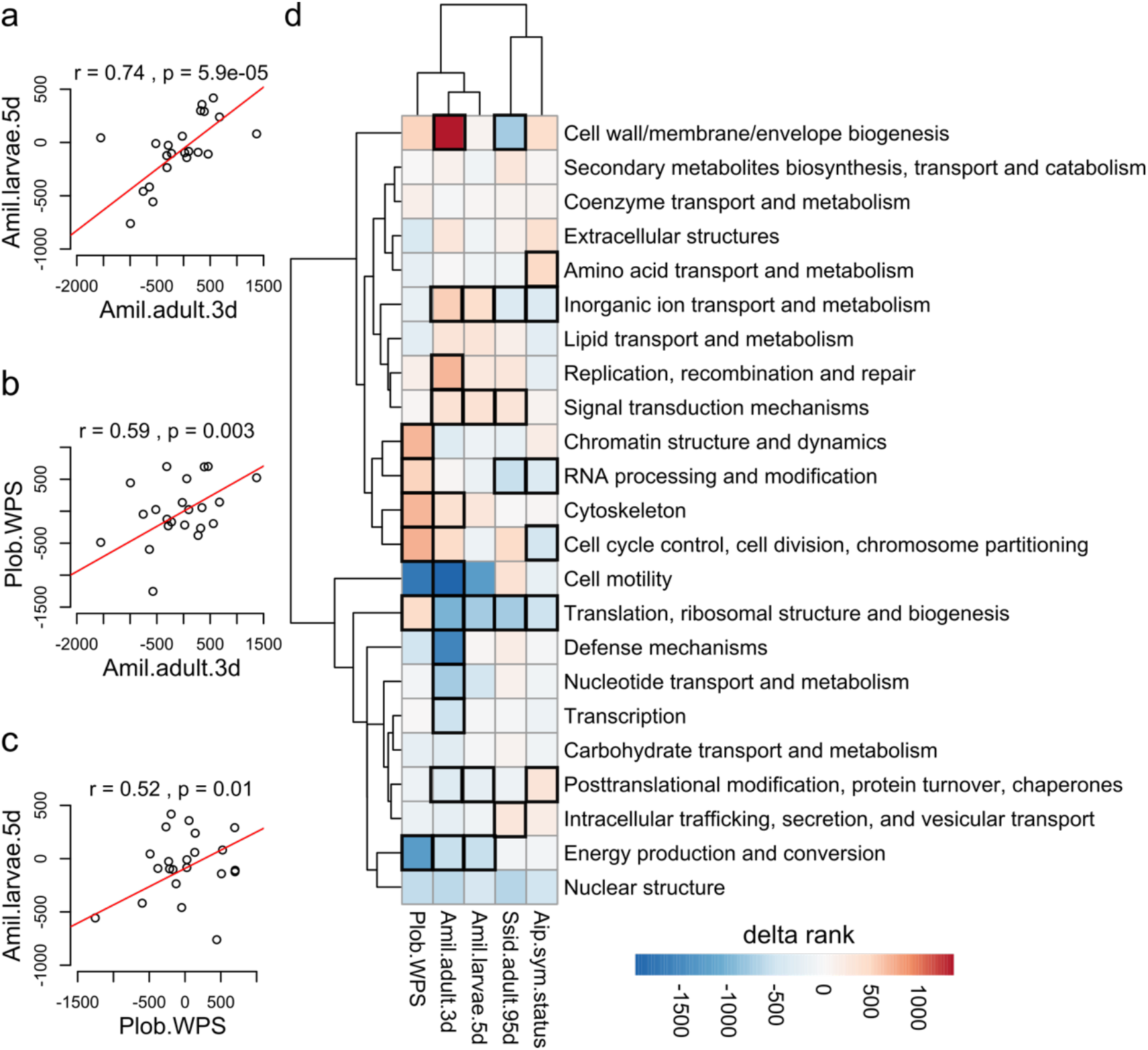
Hierarchical clustering analysis of KOG term enrichments. Correlations of KOG delta ranks between (a) *A. millepora* larvae in response to 5-day thermal stress and adults in response to 3-day thermal stress, (b) the WPS phenotype in *P. lobata* and 3-day thermal stress in adult *A. millepora*, and (c) the WPS phenotype in *P. lobata* and 5-day thermal stress in larval *A. millepora*. (d) Shared KOG term enrichments (rows) among generally upregulated or downregulated genes (delta rank heatmap) by dataset (column). Boxes outlined in bold are statistically significant enrichments (FDR-adjusted P < 0.05) within each dataset.

### Meta-analysis of expression in response to thermal stress and bleaching

Patterns of up- and down-regulation within KOG classes were used to compare WPS expression with global gene expression data generated from symbiotic adult *A. millepora* colonies in response to 3 days of heat stress (31.5°C, (Dixon et al., 2015)), aposymbiotic *A. millepora* larvae after 5 days of heat stress (31.4°C, (Meyer et al., 2011)), symbiotic adult *S. siderea* colonies after 95 days of heat stress (32°C, (Davies et al., 2016)), and menthol-bleached aposymbiotic vs. symbiotic *Aiptasia* (Matthews et al., 2017). A hierarchical clustering analysis comparing the change in magnitude and direction of differentially regulated genes within each KOG class among datasets revealed that WPS patterns were most similar to short-term stress responses in adult and larval *A. millepora* (Fig. 3). The strongest correlation was detected between *A. millepora* adults and larvae, as has been reported previously (Dixon et al., 2015) (r = 0.74, P < 0.001, Fig. 3a), but WPS expression was significantly correlated with both adult (r = 0.59, P = 0.003, Fig 3b) and larval (r = 0.52, P = 0.01, Fig 3c) patterns. No significant correlations with other datasets were detected for the *Aiptasia* or *S. siderea* datasets, which clustered distinctly from the *A. millepora* and *P. lobata* datasets (Fig 3d, S5).

Some unique patterns differentiating among the different thermal stress and bleaching datasets were also evident. Consistent significant enrichment of terms associated with replication, transcription and translation (Chromatin structure and dynamics; RNA processing and modification; Cell cycle control, cell division, chromosome partitioning; Translation, ribosomal structure and biogenesis) were evident among upregulated genes in WPS tissue and either absent or differentially regulated in other datasets (Fig. 3d). For example, ‘Translation, ribosomal structure and biogenesis’ was significantly enriched in all datasets, but the enrichment was among downregulated genes in all datasets save for WPS, where it was enriched among upregulated genes. ‘Signal transduction mechanisms’ were enriched among upregulated genes in all thermal stress datasets regardless of the duration of stress (*A. millepora* adults and larvae and *S. siderea* adults) but no enrichments were detected for the symbiotic state datasets. In symbiotic adult corals, ‘Cell wall/membrane/envelope biogenesis’ exhibited distinct patterns of regulation depending on the length of the thermal stress, being up-regulated in *A. millepora* under short-term stress, and down-regulated in *S. siderea* under long-term stress. ‘Amino acid transport and metabolism’ was enriched among upregulated genes in aposymbiotic *Aiptasia*, but not strongly regulated in any of the scleractinian coral datasets.

## Discussion

In corals impacted by WPS the loss of intracellular symbionts occurs independently of thermal stress, providing a rare opportunity to investigate the cellular and molecular mechanisms differentiating each of these processes in a reef-building coral. In spite of the clearly negative impact of WPS on holoboint physiology, we do not detect expression patterns indicative of an immune response in the coral host, suggesting that the putative viral infection resulting in WPS targets symbiont rather than host cells. Surprisingly, the meta-analysis revealed that WPS expression patterns are most similar to short-term thermal stress responses regardless of symbiotic status. While we identify some shared patterns of regulation, the striking variation among studies examining different stressor durations and symbiotic states highlights the need for additional work specifically designed to address these different processes.

The hierarchical clustering analysis of KOG term enrichments identified significant correlations between the magnitude and direction of regulation in WPS and short-term thermal stress, with patterns in *S. siderea* and *Aiptasia* being distinct and unrelated, even to each other (Fig. 3). This was unexpected because as a putative viral infection (Lawrence et al., 2015), WPS reflects a bleaching phenotype independent of thermal stress, and thermal stress responses in larval *A. millepora* are independent of symbiont status. Our initial expectation was that KOG enrichments would cluster based on stressor, with WPS being most similar to menthol bleached *Aiptasia,* and these non-thermal bleaching phenotypes distinct from the three thermal stress experiments (*A. millepora* and *S. siderea*, Fig. 3d). An alternate hypothesis was that long-term differences in symbiotic status would drive clustering, with the WPS, *S. siderea*, and *Aiptasia* datasets clustering independent of the short-term thermal stress experiments. Although it is important to point out that as we sampled corals of opportunity, we do not know when WPS symptoms first occurred in any coral in relation to our sampling time-point. Given that stress response expression patterns can be highly dynamic (Ruiz-Jones & Palumbi, 2017), future work should aim to account for sampling time to control for this potential source of variation. Nevertheless, closer examination of unique and shared patterns of differential regulation suggest that a common non-specific cellular stress response may unite the WPS and short-term stress datasets, whereas the lack of clustering among the symbiont status datasets may be driven by the difference in causative stressors or divergent host responses to a lack of symbionts.

The broad similarity in patterns of expression in WPS and short-term thermal stress may reflect non-specific cellular stress response mechanisms (Kültz, 2005), rather than those specific to bleaching. For example, down-regulation of mitochondrial proteins as observed in these three datasets (‘Energy production and conversion’, Fig 3d) is hypothesized to be a general mechanism for protecting cells from oxidative stress (Crawford, Wang, Schools, Kochheiser, & Davies, 1997). In corals, production of reactive oxygen species (ROS) increases during thermal stress and is hypothesized to overwhelm both host and symbiont detoxification mechanisms (reviewed in (C. A. Oakley & Davy, 2018; Weis, 2008). Elevated ROS production was observed in *Emiliania huxleyi* following a viral infection (Evans, Malin, Mills, & Wilson, 2006), suggesting that ROS stress may also play a role in WPS, if excess ROS are generated by symbionts during infection. Taken together, increased ROS production may help explain the similarity in global expression patterns driving clustering in these studies, although actual quantification of ROS levels in both hosts and symbionts impacted by WPS would help confirm this hypothesis.

Bleaching in WPS potentially results from a viral infection of symbionts, whereas *S. siderea* were subject to long-term exposure to elevated thermal stress (Davies et al., 2016) and *Aiptasia* were chemically bleached using menthol (Matthews et al., 2017). We did not detect significant enrichment of any immunity or stress response pathways in WPS tissues (Fig. 2). Furthermore, functional enrichment patterns are inconsistent with global expression patterns previously reported during viral infection. Inhibition of host cellular protein synthesis which conserves cellular resources and blunts host antiviral responses, is a hallmark of many viral infections (Dai et al., 2017). Whereas we observe enrichment of replication, transcription and translation among upregulated genes in the WPS dataset (Fig. 2,3). The most striking example of this difference is the consistently significant enrichment of ‘Translation, ribosomal structure and biogenesis’ which occurs among upregulated genes in the WPS dataset, but among down-regulated genes in every other dataset. Downregulation of ribosomes is a well-established response to cellular stress (López-Maury, Marguerat, & Bähler, 2008), which would be expected during thermal stress and bleaching. Yet this pattern is not evident in WPS tissues, suggesting that if a viral infection is the causative agent (Lawrence et al., 2015), it does not impact host cells. One explanation for this anomalous pattern may be that this expression reflects an effort by the host to re-establish symbiosis that is repeatedly thwarted by viral lysis of symbiont cells. An earlier study examining expression during the onset of symbiosis in other coral systems reported upregulation of genes with similar annotations, such as ‘cytoskeleton’, ‘cell cycle/growth/differentiation’ and ‘regulation of transcription’ (Voolstra, Schwarz, et al., 2009), consistent with this hypothesis; however, further work will be needed to determine the precise infection dynamics and host responses, or lack thereof.

While some similarities were evident between the *S. siderea* and *Aiptasia* datasets, for example, shared enrichment of ‘Inorganic ion transport and metabolism’ and ‘RNA processing and modification’ among downregulated genes, it is remarkable how different on average enrichment results are from one another, and from the ‘short-term’ stress response patterns. For example, the enrichment of ‘Cell wall/membrane/envelope biogenesis’ among downregulated genes is unique to *S. siderea* following 95 days of exposure to elevated temperature, whereas menthol-bleached *Aiptasia* is the only dataset exhibiting enrichment of ‘Amino acid transport and metabolism’ among upregulated genes. In *S. siderea*, expression responses likely reflect the cellular homeostasis response, rather than a cellular stress response, as would be expected in the short-term studies. The cellular homeostasis response is secondary to the immediate stress response, and acts to reestablish homeostasis under the new environmental regime (Kültz, 2005). Specific to the environmental perturbation that triggered the initial stress response, the novel patterns of regulation enacted under the homeostasis response persist until another change in environmental conditions occurs. Differential regulation enacted under the cellular homeostasis response is specific to the particular environmental variables that triggered the cellular stress response and permanent until another change in environmental conditions occurs (Kültz, 2003). Consequently these expression patterns may largely reflect physiological changes necessary to acclimatize to persistently elevated temperature, which occurs in all organisms independent of the algal symbiosis (Schulte, 2015).

Unique patterns in *Aiptasia* may reflect the metabolic flexibility of this particular symbiosis, as Symbiodiniaceae are not essential for survival. The divergence in amino acid metabolism may be a reflection of this nutritional independence, where *Aiptasia* may be more capable of regulating their metabolism to make up for a lack of symbiont nutrition than obligately symbiotic scleractinian corals. Early work in another *Aiptasia* species (*pulchella*), showed that Symbiodiniaceae synthesize and translocate seven amino acids to hosts, whereas hosts only synthesized two essential amino acids (Wang & Douglas, 1999). Loss of symbionts, and consequently, a reduction in amino acid supply, may necessitate an upregulation of the host’s reliance on heterotrophically derived nutrition, sufficient to meet daily metabolic demand (A. D. Hughes & Grottoli, 2013). As *Aiptasia* do not calcify, this upregulation may be sufficient to ensure survival in an aposymbiotic state. Although some scleractinian corals can also shift to heterotrophy during episodes of bleaching to meet metabolic demands (Grottoli, Rodrigues, & Palardy, 2006), no upregulation of genes involved in amino acid metabolism were detected in any other dataset (Fig. 3d). It is unclear if the *A. millepora* had access to supplemental food, but *S. siderea* were fed brine shrimp throughout the experimental duration (Castillo Karl D., Ries Justin B., Bruno John F., & Westfield Isaac T., 2014) and *P. lobata* were collected directly from their native reef, where polyps comprising both WPS and healthy tissue patches presumably had equal access to food. A multi-species comparison of heterotrophic feeding rates and metabolic shifts following experimental bleaching showed that *P. lobata* was unable to increase heterotrophic metabolism to compensate for symbiont loss (Grottoli et al., 2006). Nutrient stress, potentially resulting from a lack of heterotrophic feeding, was recently shown to result in a novel starvation phenotype in *Aiptasia* (Rädecker et al., 2019), consistent with the idea that heterotrophy may be key to compensating for a lack of symbionts in this system.

Taken together, we observe substantial diversity in global expression responses to thermal stress and bleaching, which may help explain the diversity of cellular bleaching mechanisms that have been reported in the literature (C. A. Oakley & Davy, 2018; Weis, 2008). As the ‘omics revolution continues, additional datasets will contribute to enhancing our understanding of the molecular and cellular processes that distinguish bleaching from thermal stress, which will be critical for understanding the mechanistic basis of the mass bleaching crisis.

### Data accessibility

Raw tag-seq reads can be obtained from NCBI’s SRA under BioProject PRJNA395362 for the WPS expression dataset. Raw reads for the *P. lobata* reference transcriptome are available at PRJNA356802. The host transcriptome assembly and associated annotation files are hosted at https://dornsife.usc.edu/labs/carlslab/data/. Bioinformatic and statistical scripts necessary to re-create analyses, as well as raw input data and annotation files used in this study are available at https://github.com/ckenkel/PoritesWPS.

## Acknowledgements

The authors are grateful for the field assistance of M Nayfa, S Noonan and J Smith. SW Davies and JL Matthews generously provided access to their raw expression data for the expression meta-analysis. CDK was supported by NSF International Postdoctoral Research Fellowship, DBI-1401165. The Australian Institute of Marine Science supported this work through the use of their research vessel (CF Ferguson) and physiological analysis through internal grants to LKB.

