## Supplementary Material for "Global gene expression patterns in response to white patch syndrome: Disentangling symbiont loss from the thermal stress response in reef-building coral"


Figure S1. Principal components analysis of physiological data. Vectors indicate loadings for each trait. Plots are identical save that individual data points are colored by (a) phenotype or (b) source colony identity (sid).

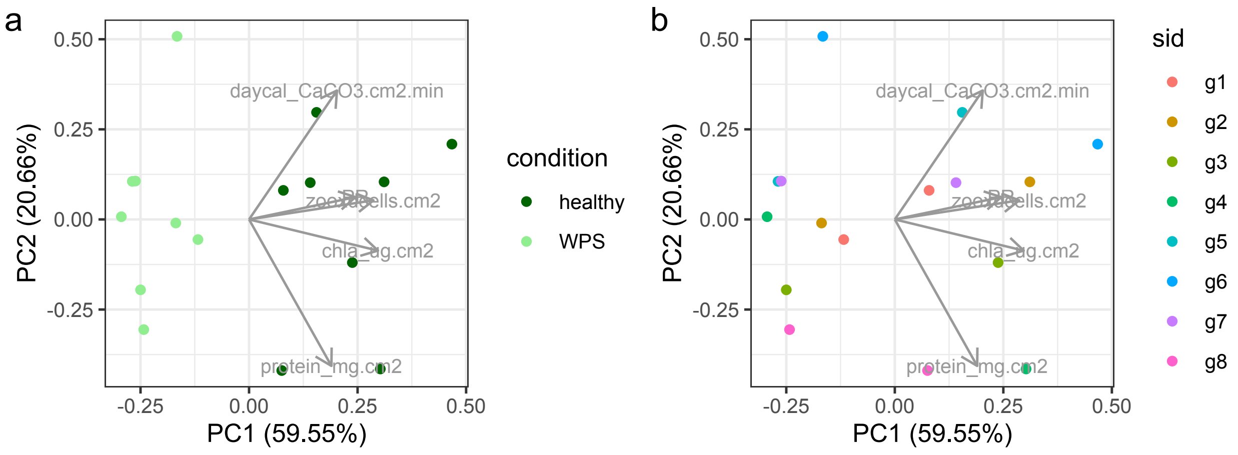


Figure S2. Venn diagram showing the number of genes differentially expressed ($P_{adj}$<0.1) by WPS phenotype (‘Patch’), source colony identity (‘Geno’), and the first principal component of the physiology dataset (‘PC1’).

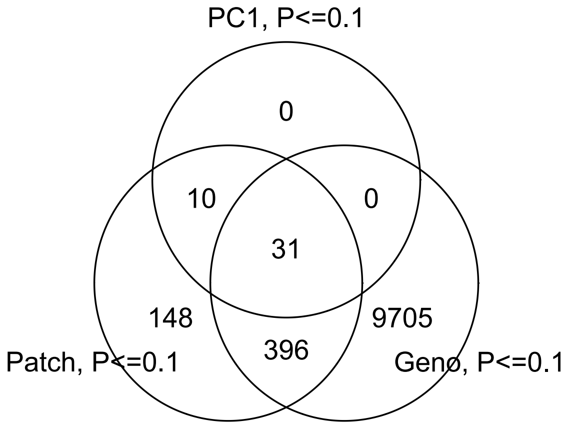

Figure S3. Hierarchical principal components analysis showing the divergence among samples for an increasing number of top differentially expressed genes by WPS phenotype (‘Patch’) and source colony identity (‘Geno’). Individual points are shaded by WPS phenotype, filled circles indicate healthy tissue while white circles indicate WPS-affected tissues.

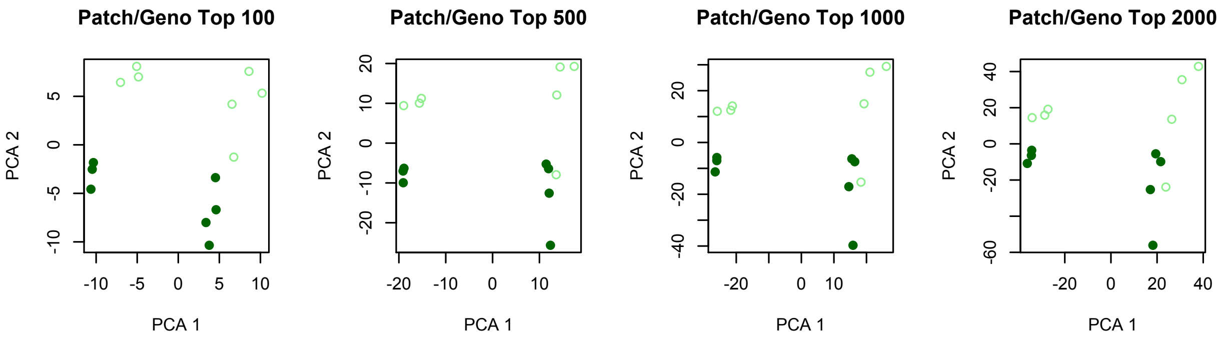


Figure S4. Heatmap of significantly differentially expressed genes (FDR-adjusted P < 0.1) annotated with the KOG term ‘Energy production and conversion’. Rows are clustered by expression similarity and columns are organized based on WPS phenotype. In column labels. Numbers indicate source colony identity, ‘B’ indicates WPS tissue and ‘N’ indicates healthy tissue.

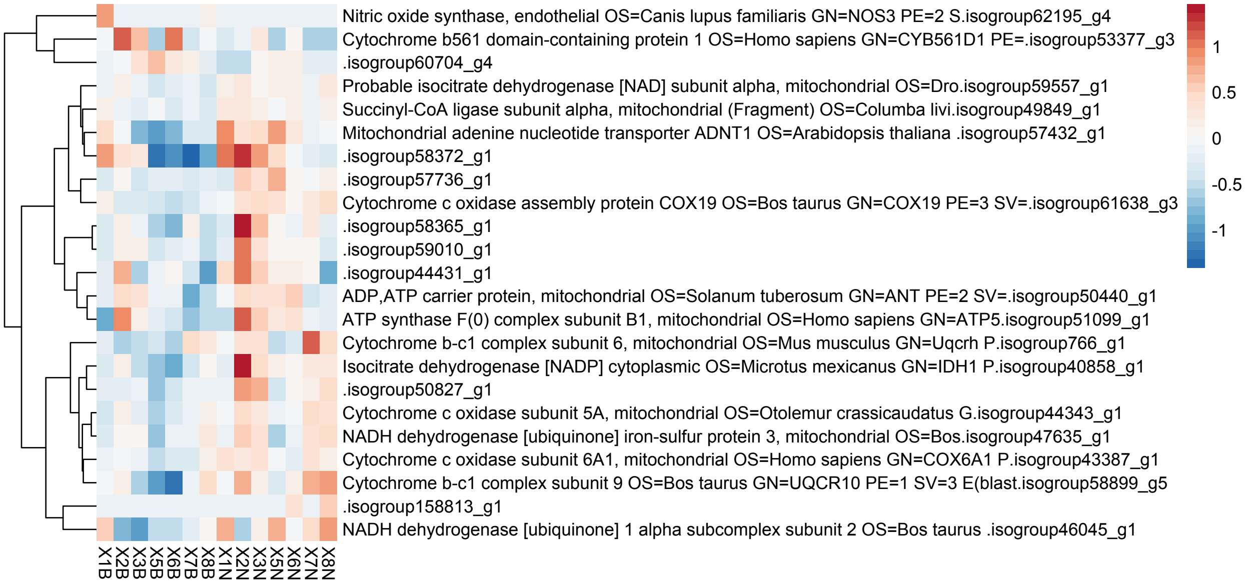

Figure S5. Pairwise Pearson correlations for KOG delta-ranks across datasets. Lower panels show individual correlations, top panels indicate the magnitude (larger font size indicates formal significance) and direction of the correlation.

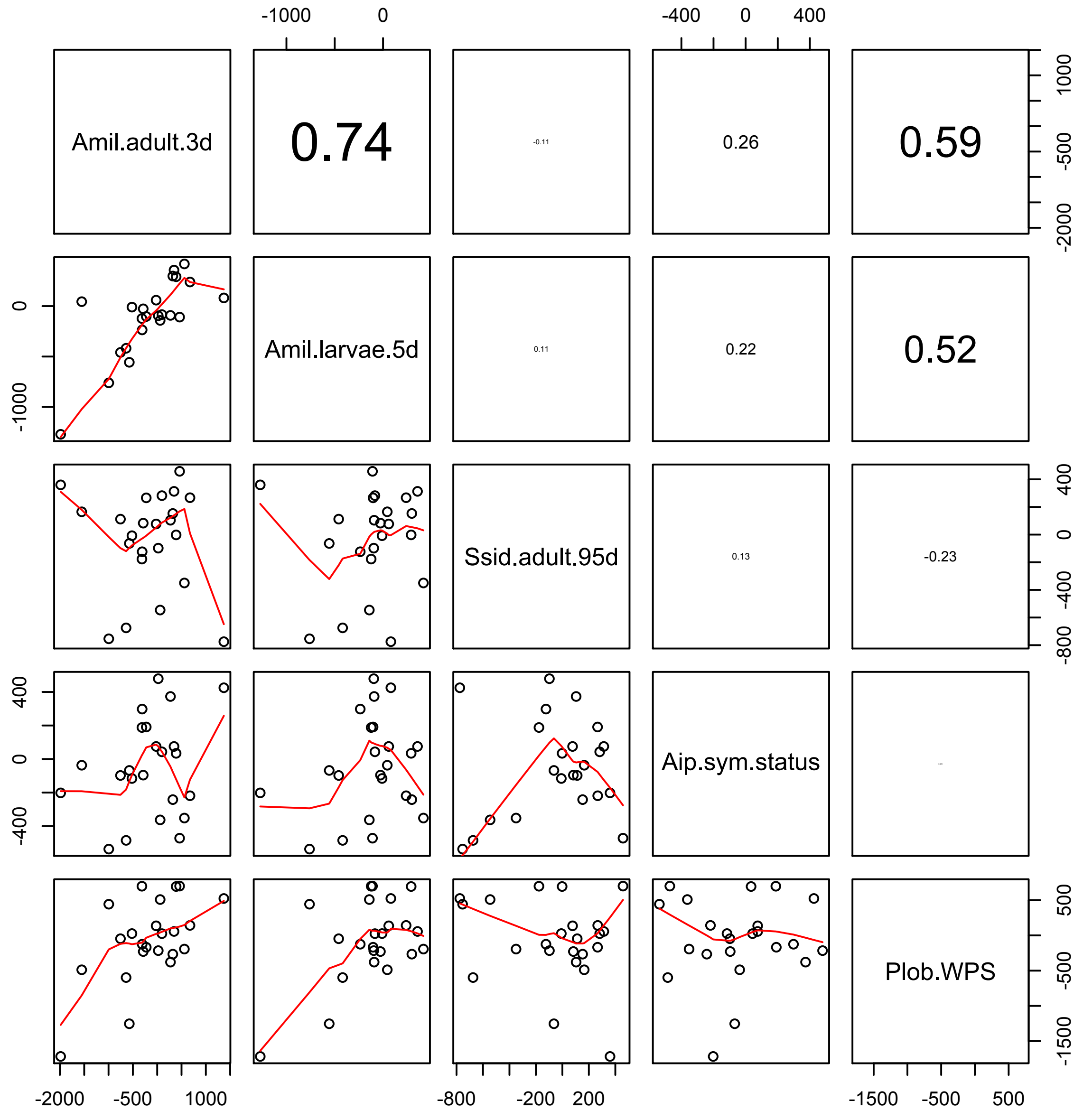
